# Senescent cells enhance newt limb regeneration by promoting muscle dedifferentiation

**DOI:** 10.1101/2022.09.01.506186

**Authors:** Hannah E. Walters, Konstantin Troyanovskiy, Maximina H. Yun

## Abstract

Salamanders are able to regenerate their entire limbs throughout lifespan, through a process that involves significant modulation of cellular plasticity. Limb regeneration is accompanied by the induction of cellular senescence, a state of irreversible cell cycle arrest associated with profound non-cell-autonomous consequences. While traditionally associated with detrimental physiological effects, here we show that senescent cells enhance newt limb regeneration. Through a lineage tracing approach, we demonstrate that senescent cells promote dedifferentiation of mature muscle tissue to generate regenerative progenitors. In a paradigm of newt myotube dedifferentiation, we uncover that senescent cells promote myotube cell cycle re-entry and reversal of muscle identity via secreted factors. Transcriptomic profiling and loss of function approaches identify the FGF-ERK signalling axis as a critical mediator of senescence-induced muscle plasticity. While chronic senescence constrains muscle regeneration in physiological mammalian contexts, we thus highlight a beneficial role for cellular senescence as an important modulator of dedifferentiation, a key mechanism for regeneration of complex structures.

## Introduction

Cellular senescence is a highly dynamic, irreversible cell cycle arrest fate induced in response to the detection of potentially genotoxic stress (Campisi, 2013). Upon senescence induction, cells permanently exit the cell cycle yet remain viable and metabolically active, exerting a strong influence over their microenvironment, most notably through the development of the Senescence-Associated Secretory Phenotype (‘SASP’ (Acosta et al., 2008; Coppé et al., 2008; Kuilman et al., 2008)). Diverse physiological events can result in senescence induction, including oncogenic transformation, developmental cues, organismal ageing and tissue injury. Nevertheless, key features of senescence including lysosomal dysfunction, resistance to apoptosis and acquisition of a secretory phenotype are largely conserved across physiological contexts and between species (Yun, Davaapil, & Brockes, 2015; Zhao et al., 2018).

Paradoxically, senescence induction can have beneficial or detrimental consequences in different contexts, depending on the nature of the senescence phenotype and the dynamics of senescent cell clearance (Walters & Yun, 2020). Cell senescence can drive tumour growth and age-related disease progression (Baker et al., 2016; Bussian et al., 2018; Childs et al., 2016; Jeon et al., 2017), yet it can also promote wound healing (Demaria et al., 2014; Jun & Lau, 2010), limit fibrosis (Kong et al., 2012; Krizhanovsky et al., 2008; Meyer, Hodwin, Ramanujam, Engelhardt, & Sarikas, 2016) and coordinate organogenesis (Davaapil, Brockes, & Yun, 2016; Muñoz-Espín et al., 2013; Storer et al., 2013). Interestingly, senescence can impact on two types of cellular plasticity of relevance to tissue renewal, namely reprogramming to pluripotency and stemness. Initially described as a cell-autonomous barrier to *in vitro* reprogramming (Banito et al., 2009), senescent cells can enhance this process in reprogrammable i4F mice by creating a tissue environment favourable to OSKM-mediated reprogramming via secreted factors (Mosteiro et al., 2016; Mosteiro, Pantoja, de Martino, & Serrano, 2018). Indeed, in the context of muscle regeneration, where a transient induction of senescence has been observed (Le Roux, Konge, Le Cam, Flamant, & Tajbakhsh, 2015), additional senescent cells derived from irradiation or ageing enhance reprogramming in an i4F background (Chiche et al., 2017). Beyond reprogramming, transient exposure to oncogene-induced SASP fosters stemness in the mammalian liver and hair follicles, though extended exposure instead promotes paracrine senescence induction, limiting tissue renewal (Ritschka et al., 2017). Further, acquisition of stem cell features by senescent cells themselves has been reported in cancer contexts and proposed to drive tissue growth (Milanovic et al., 2018). These findings suggest that cell senescence could contribute to physiological regenerative processes, and that it may modulate other types of cellular plasticity.

Dedifferentiation, a process whereby terminally differentiated cells revert to a less differentiated state within their lineage, is central to the extensive regenerative abilities found in vertebrates such as salamanders and zebrafish (Cox, Yun, & Poss, 2019; Gerber et al., 2018; Joven, Elewa, & Simon, 2019). Numerous cell types rely on dedifferentiation for the generation of regenerative progenitors in these organisms, with axolotl connective tissue (Gerber et al., 2018) and newt muscle (Wang & Simon, 2016) constituting noteworthy examples. In adult newts, limb loss triggers the formation of a blastema, a pool of lineage-restricted progenitors derived from both local stem cell activation and dedifferentiation events in mature cells from the stump, which undergoes expansion, re-differentiation and patterning to form a functional limb (Joven et al., 2019). Notably, blastema formation is accompanied by the induction of senescent cells, which are present until the onset of differentiation and are subsequently cleared by macrophages (Yun et al., 2015). Induction of senescence has subsequently been observed during zebrafish fin regeneration, where senolytic treatment slows regenerative outgrowth (Da Silva-Álvarez et al., 2020). These observations raise the possibility that cell senescence acts as a modulator of dedifferentiation, a key mechanism of appendage regeneration.

## Results

### Senescent cells accelerate blastema formation and promote myofibre dedifferentiation *in vivo*

To investigate whether and how cellular senescence impacts on regenerative processes, we leveraged a system of senescent cell induction and implantation into *N. viridescens* newt tissues (Yun et al., 2015). Limb mesenchyme-derived *N. viridescens* A1 cells were induced to undergo senescence upon DNA damage, which results in a phenotype that recapitulates many aspects of mammalian senescence including permanent cell cycle arrest, senescence-associated-β-galactosidase (SA-β-gal) activity, and SASP acquisition (Yun et al., 2015). Senescent or proliferating A1 cells were implanted into contralateral mature limb tissue of post-metamorphic newts, and the limbs subsequently amputated through the site of implantation to ensure implanted cells were present at the distal end of the remaining stump tissue (Figure 1A), the source of regenerative progenitors (Currie et al., 2016). Limbs in which control proliferating cells were implanted reached a mid-bud blastema stage three weeks post-amputation (Figure 1B). Strikingly, limbs in which senescent cells were implanted exhibited significantly larger blastema outgrowth, reaching a late-bud stage within the same period (Figure 1B-D), suggesting that senescent cells enhance blastema formation.

**Figure 1.**
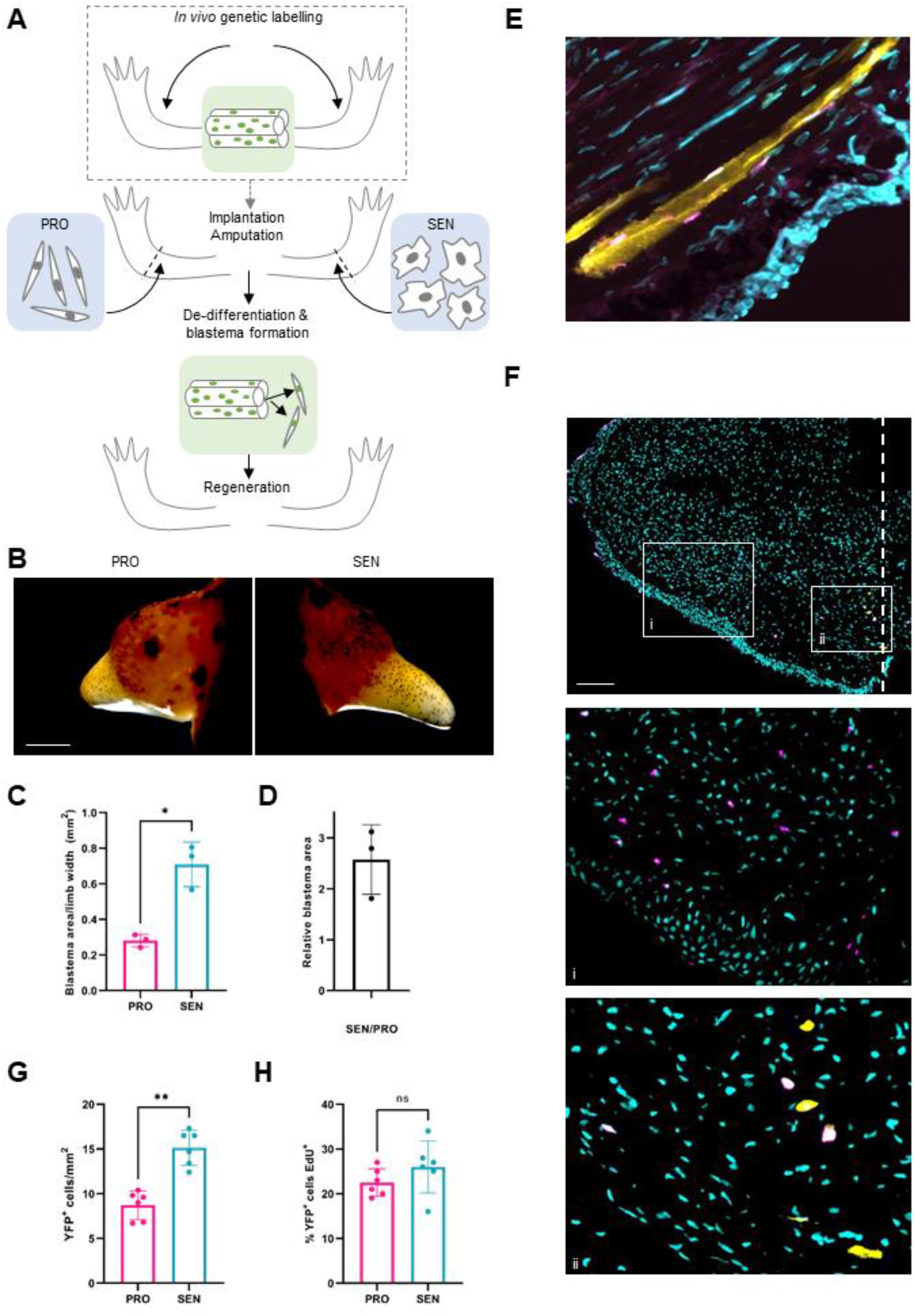
Senescent cells accelerate blastema formation and promote myofibre dedifferentiation *in vivo*. **(A)** Experimental schematic depicting fate-tracing and implantation approaches. **(B-H)** Senescent (‘SEN’) and control proliferating (‘PRO’) cells were generated *in vitro* and implanted into contralateral newt forelimbs, before amputation through the site of implantation. **(B)** Representative images of regenerating limbs at 18 days post-amputation (dpa). (*p<0.05, paired Student’s t-test, n=3). Scale bar 1000 μm. **(C)** Quantification of blastema area relative to limb width corresponding to **(B)**. **(D)** Ratio of relative blastema area from **(C)** for the indicated conditions. **(E-H)** Effect of senescent cells on myofibre dedifferentiation. Myofibres were genetically labelled (**(A)**, dashed square) prior to cell implantation and limb amputation. **(E)** Representative image of nucYFP-expressing nuclei within myofibres of the mature limb pre-amputation, as detected by α-GFP (magenta) and α-MyHC antibodies (yellow); nuclear counter-staining shown in cyan. **(F)** Representative image of an 18dpa blastema, illustrating nucYFP^+^/MyHC^+^ muscle fibres at the stump (i) and their dedifferentiated, YFP^+^/MyHC^-^ mononucleate progeny (ii). Scale bar 300 μm. **(G)** Quantification of muscle-derived dedifferentiated progenitor cells (YFP^+^/MyHC^-^) in regenerating tissue at 18dpa, after implantation of senescent or proliferating cells. **(H)** Proliferation index of YFP^+^ cells for each condition as assessed by the proportion of YFP^+^ nuclei showing EdU incorporation. **G, H**: **p<0.01, n.s. non significant, paired student t-test, n=6).

Given the importance of dedifferentiation for blastema formation, we hypothesized that senescent cells could serve as a temporary niche for promoting dedifferentiation events. We tested this notion on muscle, a tissue known to regenerate via dedifferentiation of post-mitotic myofibres in adult newts (Sandoval-Guzmán et al., 2014; Tanaka, Gann, Gates, & Brockes, 1997). Upon amputation, myofibres undergo partial loss of muscle identity, cell cycle re-entry and fragmentation, generating progenitors which proliferate, re-differentiate and fuse to form new myofibres (Wang & Simon, 2016). These progenitors can be identified through a well-established fate-tracing approach (Wang et al., 2015), where expression of a Cre recombinase under the control of a muscle-specific creatine kinase (MCK) promoter elicits nuclear YFP labelling of post-mitotic myofibre nuclei (Figure S1A). Through this approach, we genetically labelled muscle fibres in newt limbs, and subsequently repeated our implantation and amputation experiment (Figure 1A, Figure S1A). Histological analysis showed successful labelling of mature limb myofibres (YFP^+^/MyHC^+^ (Myosin Heavy Chain); Figure 1E, Figure S1B) and the appearance of YFP^+^/MyHC^-^ myofibre-derived progenitor cells in the corresponding blastema mesenchyme (Figure 1F), which have lost expression of muscle-specific genes such as myosin heavy chain (MyHC), consistent with previous observations (Wang et al., 2015). Remarkably, senescent cell implantation led to a significant expansion of the pool of dedifferentiated YFP^+^/MyHC^-^ progenitor cells (Figure 1G). The proportion of proliferating YFP^+^ cells in the blastema mesenchyme was not significantly altered (Figure 1H), indicating that the senescence-dependent increase in dedifferentiated muscle progenitors is not driven by differences in proliferation, but by the promotion of muscle dedifferentiation.

### Senescent cells promote dedifferentiation of newt myotubes through a paracrine mechanism

To further analyse the impact of senescent cells on muscle dedifferentiation, we employed an established newt myotube dedifferentiation paradigm. This system exploits the myogenic potential of the *N. viridescens* A1 cell line (Ferretti & Brockes, 1988), in which serum deprivation promotes cell cycle withdrawal and formation of multinucleate myotubes. Notably, these differentiated myotubes are able to re-enter the cell cycle upon serum exposure in culture (Tanaka, Drechsel, & Brockes, 1999; Tanaka et al., 1997; Yun, Gates, & Brockes, 2013, 2014) or upon implantation into regenerating limbs, where they contribute to the formation of muscle progenitors (Kumar, Velloso, Imokawa, & Brockes, 2000; Lo, Allen, & Brockes, 1993), modelling the muscle dedifferentiation events that occur during newt limb regeneration.

We first analyzed whether senescence was itself induced during myotube dedifferentiation, as senescence induction can occur in response to cell fusion events, such as those elicited by viruses (Chuprin et al., 2013). To this end, we induced myotube formation through serum starvation (0.25% FCS), elicited their dedifferentiation by re-exposure to 10% serum media, and analysed senescence induction based on SA-β-gal activity (Figure S2). In contrast to senescent cells, negligible SA-β-gal staining was observed in differentiated or de-differentiated myotubes (Figure S2), suggesting that senescence is not induced upon myogenic differentiation or dedifferentiation.

Next, we investigated whether senescent cells could contribute to myotube differentiation in a cell-autonomous manner, in two independent setups (Figure S3). Firstly, proliferating or senescent cells were seeded at high confluence and induced to form myotubes. Secondly, proliferating A1 cells were seeded into co-culture with senescent or control proliferating A1nGFP cells (constitutively expressing nuclear GFP (Yun et al., 2015)) under differentiation conditions. In both cases we observed that the induction of senescence coincided with the ablation of myogenic potential (Figure S3), demonstrating that senescence induction constitutes a cell-autonomous barrier to differentiation in this context.

To test whether senescent cells promote myotube dedifferentiation, as indicated by our *in vivo* data (Figure 1), we generated myotubes *in vitro,* exposed them to proliferating or senescent A1 cells and quantified the proportion of myotube nuclei undergoing cell cycle re-entry at 72h, as a readout of dedifferentiation (Figure 2A). Importantly, co-culture with senescent cells resulted in a significant increase in myotube cell cycle re-entry compared to control treatment, or fresh media alone (Figure 2B,C). The pro-plasticity effect of senescent cells was significant even in the absence of dedifferentiation-inducing conditions, suggesting that senescent cells can directly promote myotube dedifferentiation.

**Figure 2.**
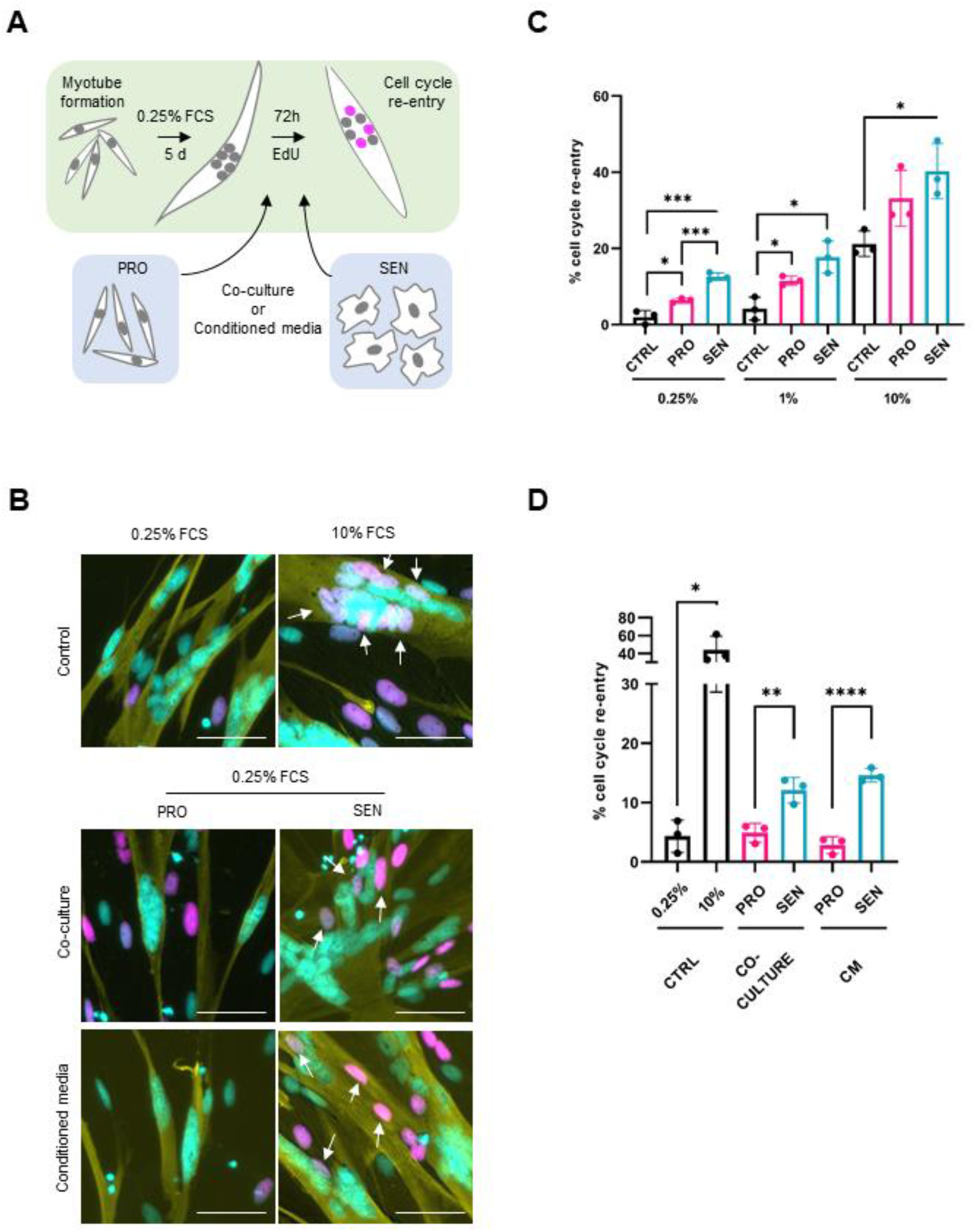
Senescent cells promote dedifferentiation of newt myotubes through a paracrine mechanism. **(A**) Schematic representation of the experimental set up. (**B**) Representative images of myotubes following immunostaining against MyHC (yellow), EdU (magenta) and Hoechst (cyan) labelling. White arrows indicate EdU^+^ nuclei within myotubes. **(C)**: Quantification of the proportion of myotube nuclei undergoing cell cycle re-entry for the indicated conditions, 72h post-treatment. PRO and SEN indicate co-culture with the respective cell populations. **(D)**: Quantification of the proportion of myotube nuclei undergoing cell cycle re-entry for the indicated conditions, 72h post-treatment. Myotubes were co-cultured or treated with conditioned media derived from the indicated populations. Two-tailed unpaired student’s t-tests were used to compare datasets (*: p<0.05, **: p<0.01, ***:p<0.001, ****: p<0.0001).

We next asked whether the viability of senescent cells is important for the observed cell cycle re-entry effects. Treatment with the senolytic agent ABT263, a BCL2 inhibitor with selective toxicity towards senescent A1 cells (Figure S4A,B), abrogates senescence-induced plasticity (Figure S4C). Interestingly, ABT263 treatment alone enhances cell cycle re-entry downstream of serum exposure, consistent with a role of apoptotic signalling in dedifferentiation (Wang et al., 2015).

We next investigated whether senescence-induced-plasticity is mediated in a paracrine manner. Treatment with senescent cell conditioned media (CM) enhances cell cycle re-entry (Figure 2B,D), recapitulating the effects of senescent cell co-culture, suggesting that senescence induced-plasticity is mediated by secreted factors. This effect is accompanied by increased phosphorylation of retinoblastoma (Rb) (Figure S5), a critical event in S-phase re-entry (Tanaka et al., 1997). Additionally, we observed a small but significant increase in proliferation of mononucleate cells upon exposure to senescent CM, suggesting that factors secreted by senescent cells exert pro-proliferative effects in mononucleate progenitor cells as well as in differentiated myotubes (Figure S6).

### Transcriptomic insights into senescence-mediated dedifferentiation

To determine the mechanistic basis for senescence-induced dedifferentiation, we performed bulk RNAseq analysis of purified cell populations (Figure 3A). We observed widespread changes to the transcriptomic profile upon the induction of senescence, differentiation or dedifferentiation (Figure 3B). Distance analysis indicated a clear segregation between sample groups and strong similarity between replicates (Figure 3C). Notably, senescent CM-treated myotubes showed greater similarity to 10% FCS-treated myotubes than those treated with fresh 0.25% media or proliferating CM (Figure 3C).

**Figure 3.**
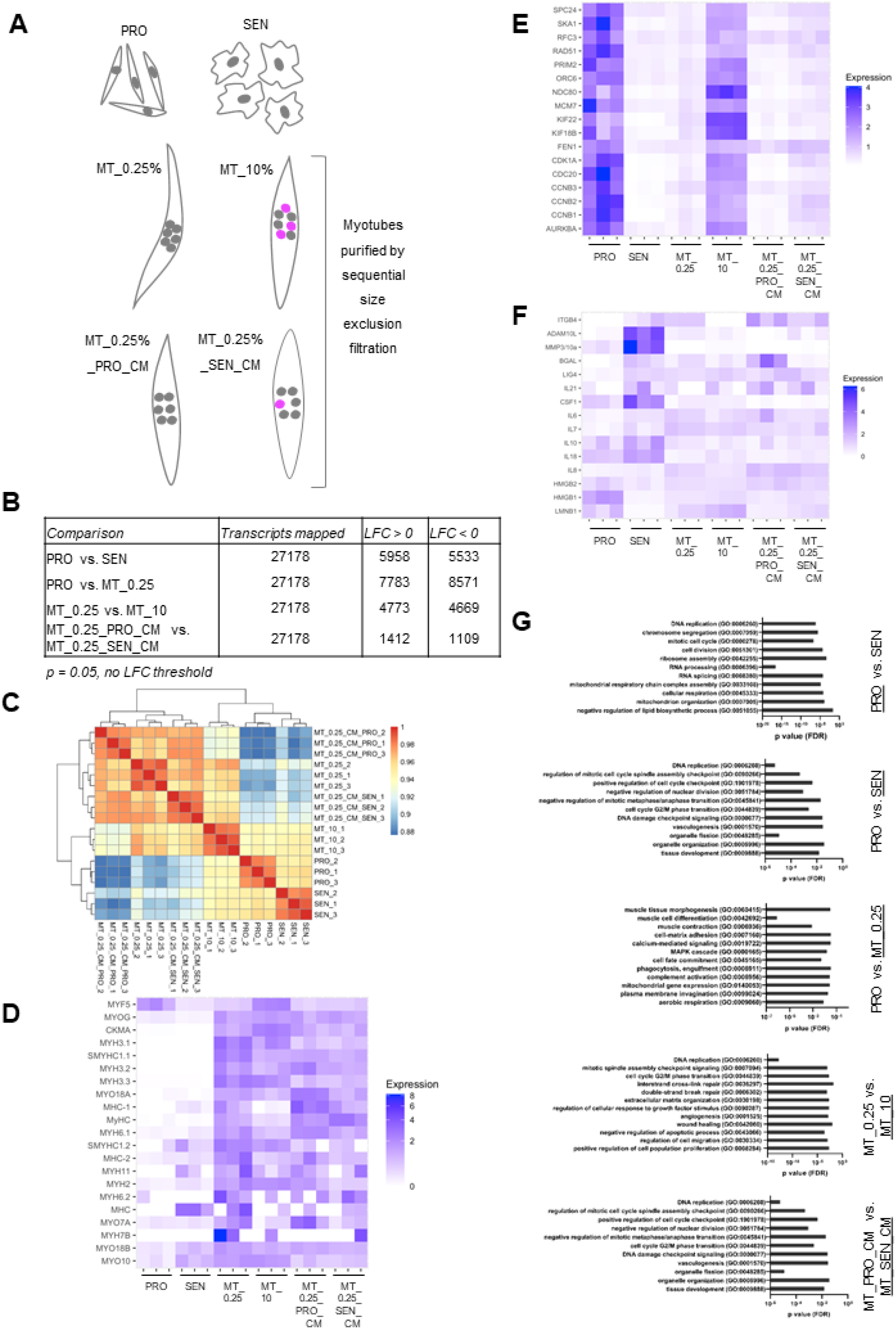
Transcriptomic insights into senescence-mediated dedifferentiation. **(A)** Experimental design schematic depicting sample groups for bulk RNAseq analysis (PRO: proliferating mononucleates, SEN: senescent cells, MT_0.25%: differentiated myotubes, MT_10%: serum-induced dedifferentiating myotubes, MT_0.25%_PROCM: myotubes cultured in proliferating cell conditioned media (0.25% FCS), MT_0.25%_SENCM: myotubes cultured in senescent cell conditioned media (0.25% FCS), all n=3). Dedifferentiating nuclei depicted in pink. **(B)** Comparison of significantly differentially regulated transcripts between sample groups. **(C)** Sample distance analysis plot. **(D-F)** Heatmaps depicting differentiation **(D**), proliferation **(E)** and senescence **(F)** related transcripts, with transcript expression for each replicate normalized relative to the mean reads per million transcripts across all sample groups. **(G)** GO-term analysis was performed using closest BLAST hits for transcripts significantly enriched (Log fold change >1, p<0.05) for the comparisons depicted (enriched population underlined).

Among the differentially regulated genes, the muscle stem cell and myoblast-associated transcription factor Myf5 showed robust expression in proliferating mononucleate cells, consistent with their myogenic potential, whereas this expression was significantly reduced in senescent cells (Figure 3D), in line with the loss of myogenic capacity upon senescence induction. Similarly, Myf5 expression was downregulated upon differentiation, coincident with increased expression of Myogenin, myosin isoforms and muscle-specific creatine kinase, indicating acquisition of differentiated myotube identity (Figure 3D). Re-exposure of differentiated myotubes to serum and, to a lesser extent, senescent cell-derived factors, reduced these readouts of differentiated muscle identity (Figure 3D).

As expected, markers of proliferation including cell cycle and DNA replication-related transcripts were significantly reduced upon induction of senescence or differentiation (Figure 3E). Of note, expression of proliferation-related transcripts was increased in serum- or senescent cell CM-treated myotubes compared to their respective controls (Figure 3E), mirroring the increased myotube cell cycle re-entry observed in both contexts (Figure 2). In the senescence compartment, we noted the conservation of important senescence-associated transcriptional changes from mammalian systems, including upregulation of classical SASP factors including IL-6, IL-8 and CSF1, as well as upregulation of several matrix-remodelling proteases, DNA repair factors and reduction in expression of nuclear architecture transcripts including HMGB1&2, and Lamin B1 (Figure 3F). GO-term analysis of significantly enriched transcripts (Figure 3G) underscored the loss of proliferative capacity upon senescence induction, where terms including ‘DNA replication’ and ‘mitotic cell cycle’ were associated with proliferating cell-enriched transcripts, and highlighted changes in RNA processing and ribosome assembly, mitochondrial and lipid metabolism upon senescence induction. Enrichment of intercellular communication-related terms (e.g. secretion, cell projection organisation), and changes related to MAPK and BMP networks were seen upon senescence induction. As expected, differentiation was associated with muscle related terms, calcium and MAPK signalling changes, while serum- and senescence-induced dedifferentiation were both associated with DNA replication, repair and cell cycle checkpoint-related terms.

### Senescent cells promote plasticity through the FGF-ERK signalling axis

To identify molecular mediators of senescence-induced plasticity, we mined our RNAseq dataset for candidates fulfilling three criteria: they should constitute secreted factors, their expression should increase upon senescence induction, and the cognate receptors and/or downstream signalling pathways for these factors should be expressed in myotubes undergoing dedifferentiation. As such, we identified several candidates belonging to the FGF, BMP, ERK and Wnt pathways, clotting factor protease activity and ECM remodelling factors (Figure S7). For each candidate pathway, we first assessed their general requirement for myotube cell cycle re-entry upon serum exposure (Figure 4A). We observed a strong suppression of cell cycle re-entry upon inhibition of BMP signalling using dorsomorphin (DMD), and MEK/ERK signalling using the inhibitor U0126 (Figure 4A), as previously observed (Wagner et al., 2017; Yun et al., 2014). Inhibition of protease activity using the broad-spectrum MMP inhibitor GM6001 or the serine protease inhibitor AEBSF, active against clotting factor protease activity (Wagner et al., 2017), had little effect on serum-induced cell cycle re-entry (Figure 4A). In contrast, inhibition of Wnt (C59) or FGFR signalling (PD173074, AZD4547) led to moderate reductions in myotube cell cycle re-entry, with the strongest effect observed using the pan-FGFR inhibitor AZD4547 (Figure 4A).

**Figure 4.**
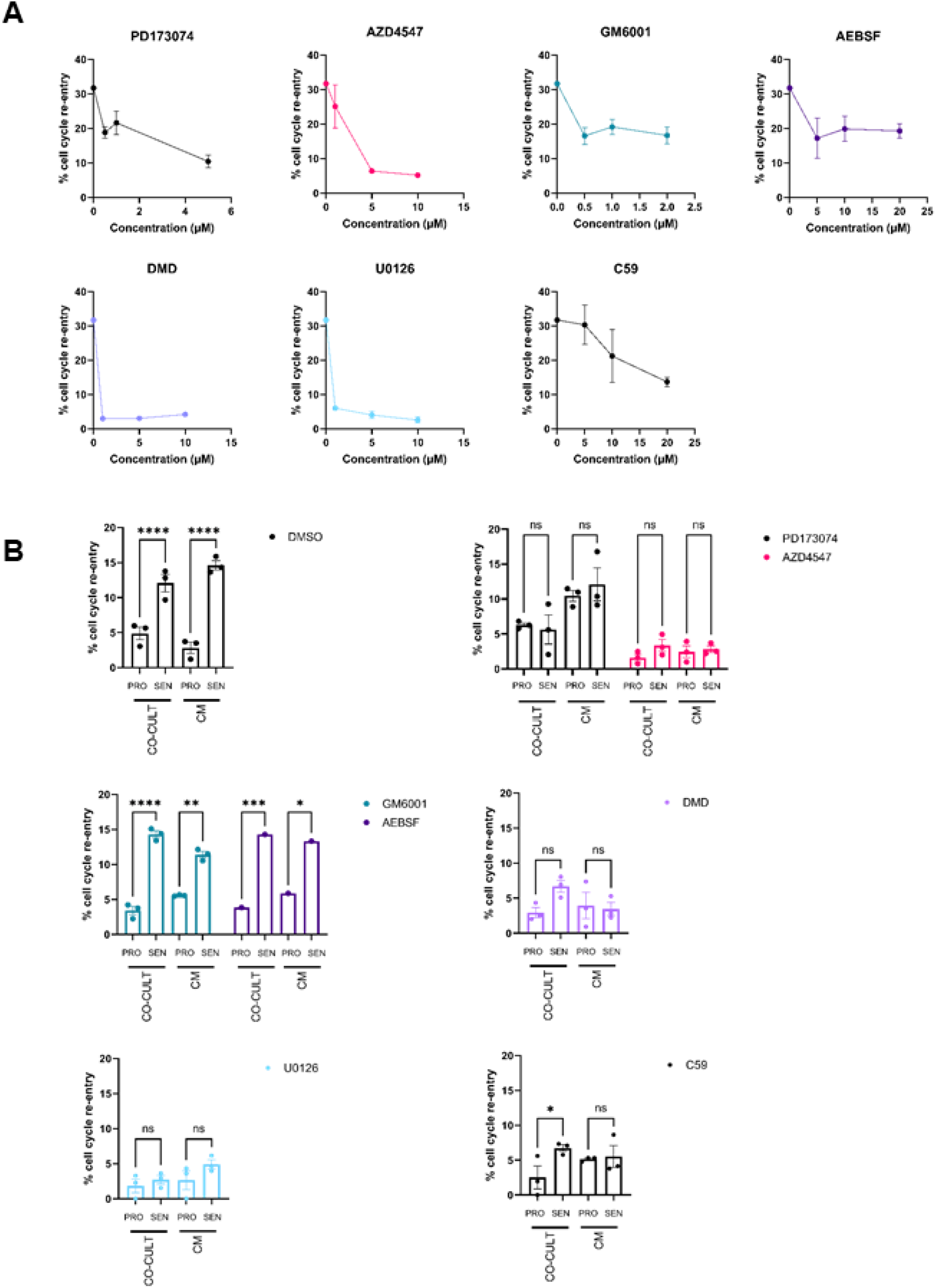
Senescent cells promote plasticity through the FGF-ERK signalling axis. **(A-B)** Quantification of the proportion of myotubes undergoing cell cycle re-rentry for the indicated conditions, 72h post-treatment. Myotubes were generated and subsequently exposed to DMSO vehicle control or inhibitors in 10% **(A)** or 0.25% **(B)** FCS in the presence of proliferating/senescent cell coculture or conditioned media treatment. In **(A)** inhibitors were used over a non-toxic dose range. In (**B)**, inhibitors were used at the following concentrations: PD173074 1 μM, AZD4547 5 μM, GM6001 2 μM, AEBSF 20 μM, U0126 10 μM, DMD 1 μM, C59 10 μM. Cell cycle re-entry was quantified as the proportion of myotube (MyHC^+^) nuclei showing EdU incorporation. Statistical analysis was performed using and Tukey post-hoc multiple comparisons testing (*: p<0.05, **: p<0.01, ***: p<0.001, ****: p<0.0001, ns: not significant). Representative data from n>2 experiments shown.

We next tested the importance of these factors for senescence-induced plasticity (Figure 4B). Blocking BMP or ERK signalling abrogated serum-induced cell cycle re-entry (Figure 5B). Senescence-induced plasticity is likely to rely on ERK signalling within de-differentiating myotubes to coordinate cell cycle re-entry, in light of previous work (Yun et al., 2014). Further, soluble BMP factors have been shown to promote cell cycle re-entry (Wagner et al., 2017), though intriguingly, we observed notably larger myotubes following DMD treatment, suggesting a possible role for the inhibition of BMP signalling in facilitating myogenic differentiation. Indeed, we observed increased myogenesis of A1 cells in high serum medium upon treatment with 1 μM DMD (Figure S8). These data suggest an additional role for BMP signalling in myogenic differentiation, and are consistent with previous reports examining BMP inhibition in C2C12 murine *in vitro* myogenesis (Horbelt et al., 2015). It is thus likely that BMP inhibition promotes differentiation and blocks dedifferentiation irrespective of stimuli derived from serum or senescent cells.

While protease inhibition had no effect on senescence-induced plasticity (Figure 4B), inhibition of Wnt signalling decreased the effect of senescent cell co-culture and abrogated that of senescent CM (Figure 4B), suggesting that WNT ligands contribute to this process. Further, inhibiting FGFR signalling using PD173074 (FGFR1) or AZD4547 (FGFR1-3) resulted in blockage of senescence-induced plasticity by co-culture or CM, suggesting a critical role of FGF signalling in mediating this process (Figure 4B). Given the upregulation of secreted FGF ligands in senescent cells, the robust expression of FGF receptors in de-differentiating myotubes (Figure S7), as well as the critical role of the FGF-effector ERK in cell cycle re-entry, the FGF-ERK axis emerges as a direct mediator of senescence-induced plasticity (Figure S9).

## Discussion

This study demonstrates that cellular senescence can play beneficial roles during salamander limb regeneration. Specifically, we show that senescent cells promote muscle dedifferentiation, a critical process underlying successful limb regeneration, and uncover that they are able to modulate muscle plasticity directly, through the secretion of paracrine factors including WNT and FGF ligands.

As such, our findings provide important advances for our understanding of senescent cell functions as well as early events during limb regeneration, opening up several research avenues. With regards to limb regeneration, the development of *in vivo* senescent cell labelling and depletion approaches in newt species should enable in-depth explorations of the physiological functions of these significant cellular players in the future. In addition, as dedifferentiation has been shown to underlie axolotl connective tissue regeneration (Gerber et al., 2018), it would be of interest to assess if senescent cells modulate this process. Further, probing the impact of senescent cells on transdifferentiation, as it happens in the salamander lens (Tsonis, 2006), will be relevant for understanding their roles in different plasticity contexts.

It remains possible that senescent cells play additional roles during salamander limb regeneration. Indeed, our data suggests that these cells can promote cell proliferation (Figure S6). While we have not observed pro-proliferative effects specific to the muscle progenitors (Figure 1H), it remains possible that senescent cells promote proliferation of additional blastema populations, which may explain the notable acceleration of blastema development observed upon senescent cell implantation (Figure 1B-D). This is in agreement with unpublished data from our group (Yu et al, in submission), which reports that senescent cells facilitate progenitor cell expansion in the axolotl. Additionally, senescent cell clearance in salamanders is achieved by macrophages (Yun et al., 2015), raising the possibility that senescent cells may have indirect functions via recruitment or regulation of immune cell activity, important in other regenerative contexts (Ratnayake et al., 2021). Lastly, the immune-dependent clearance mechanism acting in salamanders may be critical to ensure that senescence induction has beneficial rather than detrimental consequences, limiting cell senescence to a short time-window following injury and thus creating a transient niche permissive to dedifferentiation.

FGF signalling stands out as a key mediator of senescence-induced dedifferentiation (Figure 4). In agreement, chemical approaches suggest FGFR1 activity is central for dorsal iris pigment epithelial cell dedifferentiation during newt lens regeneration (Del Rio-Tsonis, Trombley, McMahon, & Tsonis, 1998), and for blastema formation in the zebrafish fin (Poss et al., 2000). Similarly, the coordinated activity of FGF and BMP signalling - originating from the dorsal root ganglia – contributes to blastema formation in axolotl limb regeneration (Satoh, Makanae, Nishimoto, & Mitogawa, 2016). Highlighting the evolutionary conservation of pro-regenerative roles of FGF ligands, FGF4 promotes limb outgrowth in chicken (Kostakopoulou, Vogel, Brickell, & Tickle, 1996), while FGF2 is upregulated during blastema growth in murine digit tip regeneration (Takeo et al., 2013). Further careful analysis will be required to elucidate which of the senescence-upregulated newt FGF ligands is responsible for the promotion of cellular plasticity in this context. Additionally, our work uncovers WNT factors as contributors to the plasticity effect. How WNT and FGF pathways interact to promote dedifferentiation warrants further investigation.

Together, our findings uncover a beneficial role for cellular senescence during newt limb regeneration through the non-cell-autonomous promotion of muscle plasticity. In contrast, chronic senescence constrains muscle regeneration in physiological mammalian contexts (García-Prat et al., 2016). In light of our data, examination of cross-species differences in senescent cell nature, senescent-progenitor and immune crosstalk at the site of injury, and dynamics of senescence induction and clearance, could be instructive for limiting the deleterious effects of senescent cells in mammals and harnessing their beneficial traits in clinical settings.

## Supporting information

Supplemental material combined

## Acknowledgments

We thank Heng Wang for the donation of lineage-tracing plasmids, the Dresden Concept Genome Centre (CRTD, TU Dresden) for performing RNAseq, Phillip Gates for technical support, and all members of Yun lab for advice and comments on the manuscript. HEW was supported by an Alexander von Humboldt postdoctoral research fellowship and a TU Dresden Graduate Academy ‘Postdoc Starter Kit’ grant. KT is supported by a DAAD Scholarship. MHY is supported by Deutsche Forschungsgemeinschaft grants (DFG 22137416 & 450807335 & 497658823) and TUD-CRTD core funds.

## Author contributions

HEW, KT and MHY designed and performed experiments, analysed and interpreted data. HEW wrote the manuscript with input from all authors. HEW and MHY acquired funding. MHY supervised the study.

## Data availability

Data pertaining to this study is included within this manuscript and its supplementary materials are available upon request from MHY.

## Methods

### Animal husbandry

Axolotls (*A. mexicanum*) were obtained from Neil Hardy Aquatica (Croydon, UK) and from the axolotl facility at TUD-CRTD Center for Regenerative Therapies Dresden (Dresden, Germany). Animals were maintained in individual aquaria at ~18–20 °C, as previously described (Oliveira et al., 2022). Axolotls of the leucistic (d/d) strain were used in all experiments. All animal procedures used in this study were performed in compliance with the Animals-Scientific Procedures-Act 1986 (United Kingdom Home Office), and the laws and regulations of the State of Saxony, Germany.

### Animal procedures

Procedures for care and manipulation of *N. viridescens* newts used in this study were carried out in compliance with the Animals (Scientific Procedures) Act 1986, approved by the United Kingdom Home Office.

Tracing of de-differentiated progenitor cells was performed as described (Wang et al., 2015) with the following modifications: plasmids (MCK:Cre, CMV:Tol2-transposase; CAG:loxp-cherry-stop-loxp-h2bYFP) were purified by Caesium chloride preparation. Electroporations were carried out using a SD9 Stimulator device as previously described (Yun et al., 2013).

For cell implantation, senescent and control proliferating cells were generated as below. Newts were anaesthetised in 0.1% tricaine and 2,000 cells were subsequently implanted into contralateral limbs using 10 μl Hamilton syringe (Hamilton, Reno, NV) with a 30^1/2^ g, 45° tip needle (Hamilton) attached to a micromanipulator, using Fast Green to track the distribution of cell solution within the tissues, as described (Yun et al., 2015). Implantations were carried out under a Zeiss Axiozoom V.16 fluorescence stereomicroscope and cells were implanted along electroporated area based on mCherry fluorescence. Newts were amputated at the mid-humerus level through the site of cell implantation/tissue electroporation under a Zeiss Axiozoom V.16 fluorescence stereomicroscope. Animals were allowed to regenerate at 20 °C. To detect EdU incorporation, 10 mM EdU (20 μl per animal) were administered by intraperitoneal injection. After 96 hr, tissues were collected and processed as described below.

### Tissue sectioning and histology

For analysis of dedifferentiation, regenerating limbs were collected by amputation and fixed in 4% (wt/vol) ice-cold paraformaldehyde (PFA) for 16–18 h at 4 °C, washed twice in PBS, and embedded in Tissue Tek-II. The samples were sectioned longitudinally in a cryostat at 10 μm. Sections were collected in Superfrost slides and stored at −20 °C. Antibody staining of tissue sections was performed using standard protocols with the indicated antibodies (Table 2). EdU incorporation was determined subsequent to immune-staining with anti-YFP antibodies using Click-iT Edu Alexa Fluor 594 Imaging kit (Life Technologies).

### Cell culture

*N. viridescens* limb-derived A1 cells (Ferretti & Brockes, 1988) and A1n*gfp* cells (Yun et al., 2015) were cultured as previously described (Yu et al., 2022); in brief, cells were grown on gelatin-coated flasks in MEM (Gibco) supplemented with 2 nM L-glutamine (Gibco), 10 μg/ml insulin (Sigma), 100 U/ml penicillin/streptomycin (Gibco), 10% heat-inactivated FCS (Gibco) and 25% v/v dH_2_O. Cells were passaged 1:2 when approaching 70-80% confluence and maintained at 25°C and 2% CO_2_.

### Induction of differentiation and dedifferentiation

A1 cells were used to generate myotubes. Cells were seeded into gelatin-coated wells at high density (2×10^4^ cells/cm^2^) and subsequently cultured for 5 days in culture media with 0.25% FCS to promote differentiation. To assay cell cycle re-entry, cultures were then exposed to fresh media for 72 hours (supplemented with 0.25, 1% or 10% FCS (PAA) as described) containing 5 μM EdU, or with co-culture and conditioned media treatment as described.

### Senescence induction and conditioned media generation

*In vitro* senescence induction was performed according to Yu, Walters and Yun (2022), using UV-irradiation (3J/m^2^, UV Stratalinker) or 24-hour exposure to 20 μM etoposide, both followed by treatment with 1 μM Nutlin-3a. Senescence induction following 12 days’ treatment was confirmed by positive SA-β-gal staining, a significant reduction in EdU incorporation, expansion of mitochondrial and lysosomal networks and persistent DNA damage foci (Yu et al., 2022). Control proliferating cells were seeded in parallel, treated only with identical volumes of DMSO, and passaged during the 12-day senescence induction to avoid confluence. For co-culture assays, proliferating and senescent cells were harvested, cells were counted using an automated cell counter (Scepter 2.0, Millipore) and seeded into co-culture at a 1:10 ratio to cells initially seeded for myogenesis (i.e. 2×10^3^ cells/cm^2^), mimicking the proportion of senescent cells reported in blastemas *in vivo* (Yun et al., 2015). For conditioned media treatment, 10 cm plates of proliferating or senescent cells at comparable confluence were washed in 80% PBS (‘A-PBS’), before incubation with 10 ml fresh media for 48 hours. Conditioned media was subsequently collected and passed through a 0.22 μm filter prior to use. Fresh conditioned media was used in every experiment.

### Inhibitor treatments

Drug toxicity assessment was performed by seeding cells into 96 well plates before subsequent treatment with a dose curve of each inhibitor for 72 hours. Cell viability was then assessed using the alamarBlue assay according to manufacturer’s instructions. For small molecules used in dedifferentiation assays, drug doses selected maintained >80% of control cell viability by alamarBlue assessment (data not shown). For ABT263 senolytic assessment, control proliferating and senescent cells were assessed in parallel, and toxicity was also assessed by dead cell staining with TO-PRO-3 iodide (Invitrogen), following incubation at 1 μM for 30 minutes in in fresh media together with NucBlue Live (Invitrogen) and microscopy assessment.

### EdU, SA-β-gal and immunostaining

SA-β-gal staining was performed according to manufacturer’s instructions (Cell Signalling) as described (Yun et al., 2015), prior to permeabilisation, EdU or immuno-staining procedures. Click-iT EdU staining was performed according to manufacturer’s instructions (Invitrogen); in brief, cultures were fixed in 4% PFA at 4°C for 10-15 minutes, permeabilised with 0.2% Triton-X100 in PBS and stained with the Click-iT reaction cocktail. Prior to immunostaining, samples were blocked in 10% goat serum in PBS for >30 minutes (RT), and immunostaining was performed (see Table 2 for details of antibodies used). Primary antibodies were incubated overnight at 4°C and secondary antibodies for 1-4 h at room temperature. Antibodies were diluted in 5% goat serum and 0.1% Triton-X100 in PBS, and samples were washed twice in PBS between primary and secondary incubation. Nuclei were counterstained using Hoechst 33342.

### Imaging

Imaging of *in vitro* fluorescence experiments was performed using a Nikon Eclispe TsER microscope. A Zeiss AxioZoom V16 microscope was used to perform SA-β-gal imaging. FIJI was used for image analysis. Imaging of *in vivo* dedifferentiation experiments was conducted using a Zeiss AxioZoom V.16 fluorescence stereomicroscope and Zen software (Zeiss). For each sample, 10 sections were scored. Blind counting was employed for all quantifications.

### Western blotting

For Western blotting, myotube cultures were lysed *in situ* using 0.02 M Hepes (pH 7.9), 0.2 mM EDTA, 1.5 mM MgCl2, 0.42 M NaCl, 25% glycerol lysis buffer, incubated for 30 min at 4°C, and subsequently cleared of debris by centrifugation. Protein concentration was analysed by the Bradford assay, and denatured lysates of equal protein amount were loaded onto 10% Bis-Tris Novex gels and run at 150V for 1h in MOPS-SDS running buffer. Overnight transfer onto nitrocellulose membranes was performed in methanol transfer buffer, before membranes were blocked (Odyssey blocking buffer, Licor, 30 minutes RT) and probed with primary antibodies in blocking buffer (>1h), before washing in PBS-T and probing with secondary antibodies. Blots were thoroughly washed in PBS and scanned using an Odyssey scanner (Licor). Bands were quantified from triplicate samples against loading controls using FIJI.

### RNAseq

For myotube purification, cultures were washed gently with A-PBS, lifted using trypsin, then quenched using fresh media. Cell suspensions were passed sequentially through 100 μm filters (to exclude aggregates) and 35 μm filters. Myotubes retained on the 35 μm filters were collected in fresh media and spun down. Proliferating and senescent mononucleate cultures were not filtered, but were simply lifted and spun down. Myotube or cell pellets were immediately lysed in buffer RLT and RNA extracted using RNeasy mini (mononucleate) or micro (myotube) kits according to manufacturer’s instructions. cDNA synthesis and RNA sequencing was subsequently performed by the Dresden Concept Genome Center (DCGC). For bioinformatic analysis, useGalaxy.org (Afgan et al., 2018) was used for initial processing. Firstly, adapter sequences were trimmed from FASTQ files using TrimGalore, quality control performed using FastQC. Alignment of trimmed reads to the *N. viridescens* transcriptome (Abdullayev, Kirkham, Björklund, Simon, & Sandberg, 2013) was performed using Sailfish. Data were then imported into Rstudio for normalization and differential gene expression analysis using DESeq2 with a significance cutoff of p<0.05.

### Statistical analysis

Statistical analysis was performed using Prism software; for comparison of n>2 sample groups, ANOVA and post-hoc Dunnett or Tukey tests were performed, and for comparison of n=2 sample groups, two-tailed student t-tests were applied as described.

