## Supplemental material combined for "Senescent cells enhance newt limb regeneration by promoting muscle dedifferentiation"

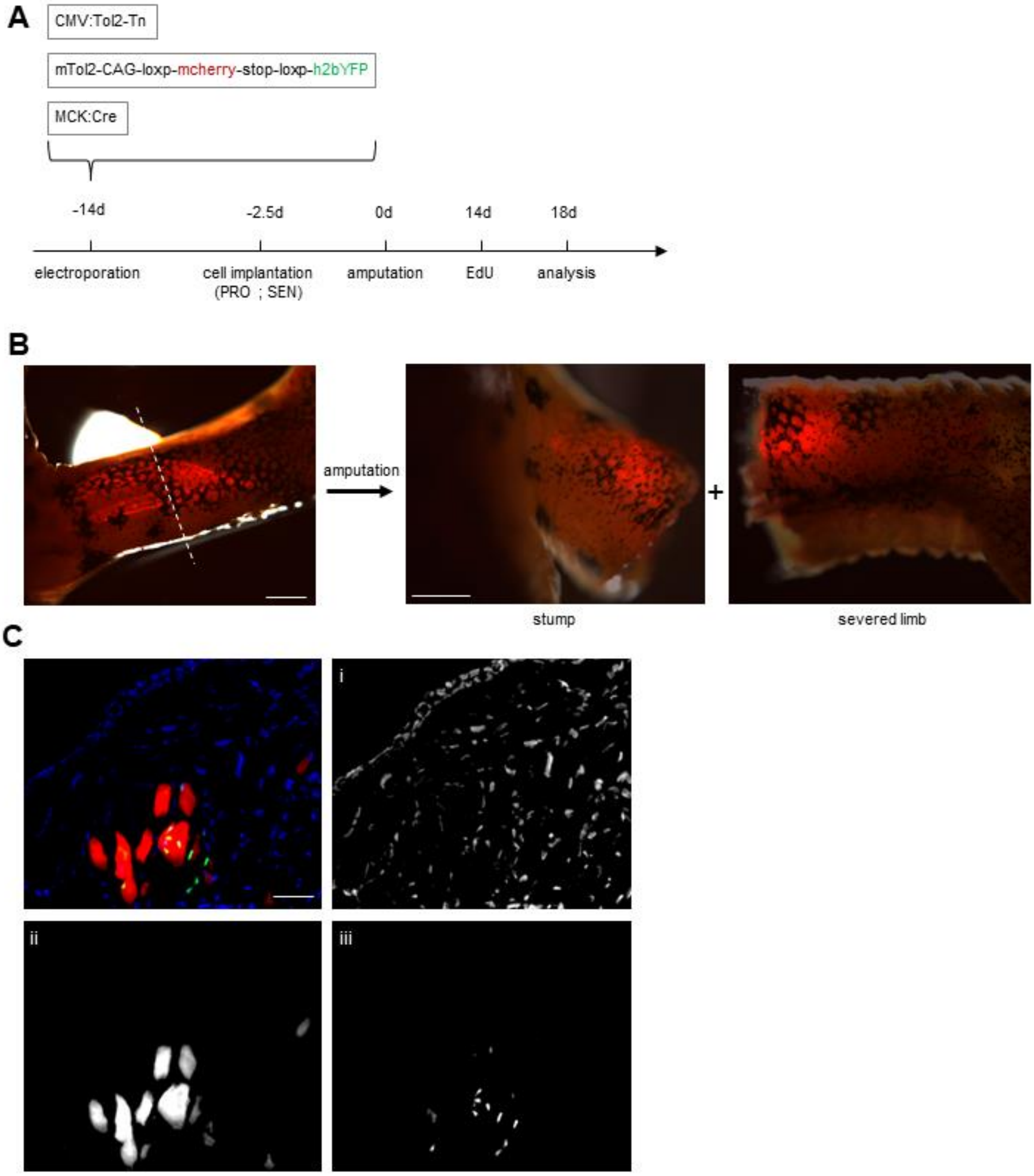
A
CMV:Tol2-Tn
mTol2-CAG-loxp-mcherry-stop-loxp-h2bYFP
MCK:Cre
-14d
-2.5d
0d
14d
18d
electroporation
cell implantation
(PRO ; SEN)
amputation
EdU
analysis
B
amputation
+
stump
severed limb
C
i
ii
iii

**Supplementary Figure 1. Genetic strategy for labelling myofibres and fate-tracing dedifferentiated progenitors *in vivo*.**

**(A)** Schematic depicting *in vivo* myofibre labelling approach to enable fate-tracing of muscle-derived dedifferentiated progenitor cells during regeneration.

**(B)** Representative image of a newt limb following electroporation with the indicated constructs and cell implantation (left, at 0d). Limb amputation, performed through the centre of the electroporated area (yellow dotted line), results in a stump (middle) which contains genetically labelled myofibres. Scale bar 500 μm.

**(C)** representative immunofluorescence image of a stump cryosection depicting $YFP^{+}$ /$MyHC^{+}$ myofibres, as detected through α-GFP (green, iii) and α-MyHC (red, ii) antibody staining, with nuclear counterstaining shown in blue (i). Their progeny can be subsequently traced during limb regeneration through the expression of nuclear YFP. Scale bar 50 μm.

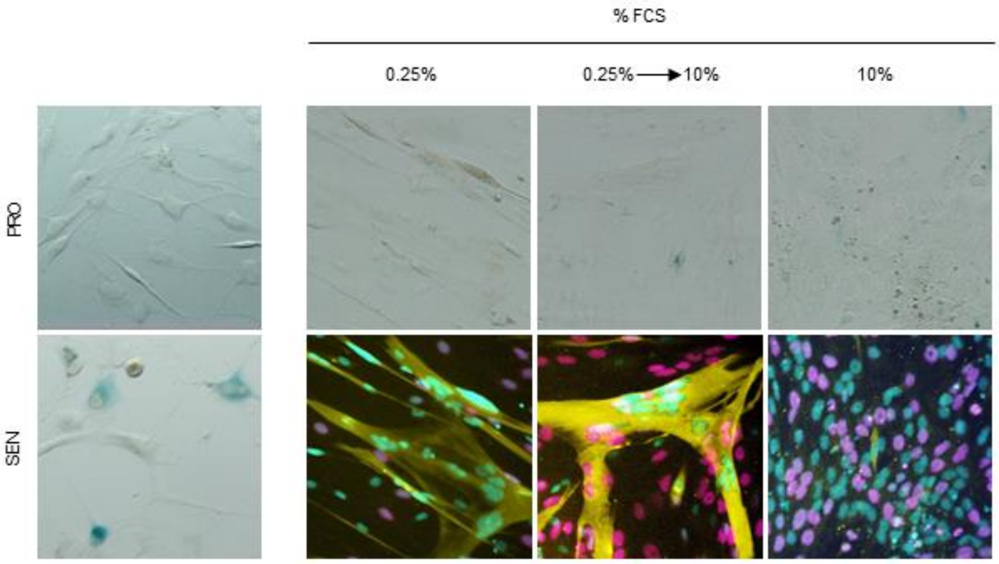
% FCS
0.25%
0.25% → 10%
10%
PRO
SEN

**Supplementary Figure 2. Senescence is not induced by cell-cell myogenic fusion or dedifferentiation events.**

Top row: Representative images of differentiated (0.25% FCS), dedifferentiated (0.25% → 10% FCS) or undifferentiated A1 cultures (10% FCS) after SA-β-gal staining (blue in brightfield). Control senescent (‘SEN’) and untreated proliferating (‘PRO’) cultures were used as positive and negative controls respectively.

Bottom row: Representative images of the indicated cultures after staining against α-MyHC (yellow), EdU (magenta) and Hoechst (cyan). Note EdU incorporation in dedifferentiating myotube nuclei (magenta). n=4.

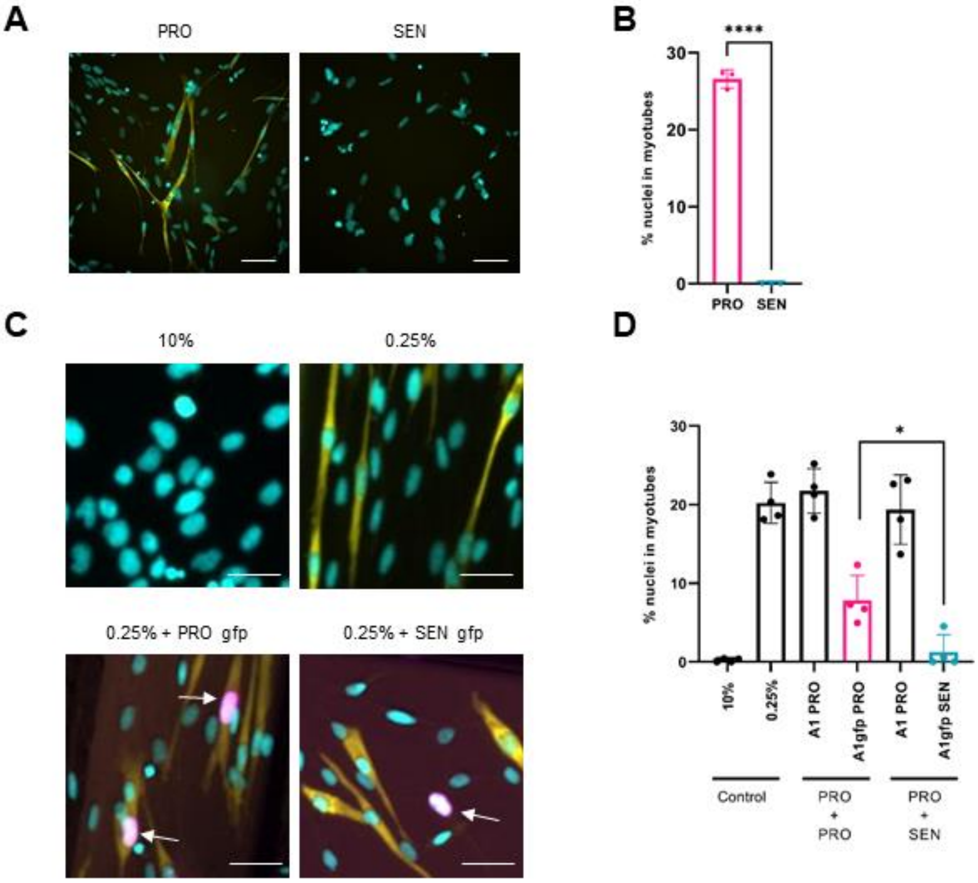

A
PRO
SEN
B
****
% nuclei in myotubes
30
20
10
0
PRO
SEN
C
10%
0.25%
0.25% + PRO gfp
0.25% + SEN gfp
D
*
% nuclei in myotubes
30
20
10
0
10%
0.25%
A1 PRO
A1gfp PRO
A1 PRO
A1gfp SEN
Control
PRO + PRO
PRO + SEN

**Supplementary Figure 3. Senescence induction ablates myogenic potential.**

**(A, B)** Representative images of senescent (SEN) and control proliferating (PRO) cells, 5 days after treatment with differentiation media (0.25% FCS) and subsequent immunostaining (Hoechst; cyan, α-MyHC; yellow). The proportion of nuclei present in myotubes was quantified **(B)** from n>50 nuclei per replicate (n=3).

**(C)** Representative images of cultured cells following immunostaining against MyHC (yellow) and GFP (magenta), and counterstaining with Hoechst (cyan). A1n*gfp* senescent ('SEN gfp') and control proliferating ('PRO gfp') cells were generated *in vitro* and subsequently lifted and seeded into co-culture with A1 cells at high confluence. Co-cultures were then treated with differentiation media for 5 days before fixation and staining. White arrowheads indicate $GFP^+$ nuclei.

**(D)** Percentage of nuclei present in myotubes (n>50 nuclei per replicate, n=3).

Two-tailed unpaired student's t-tests were used to compare myogenesis between proliferating and senescent cells in both experimental set-ups (*: p<0.05, ****: p<0.0001).

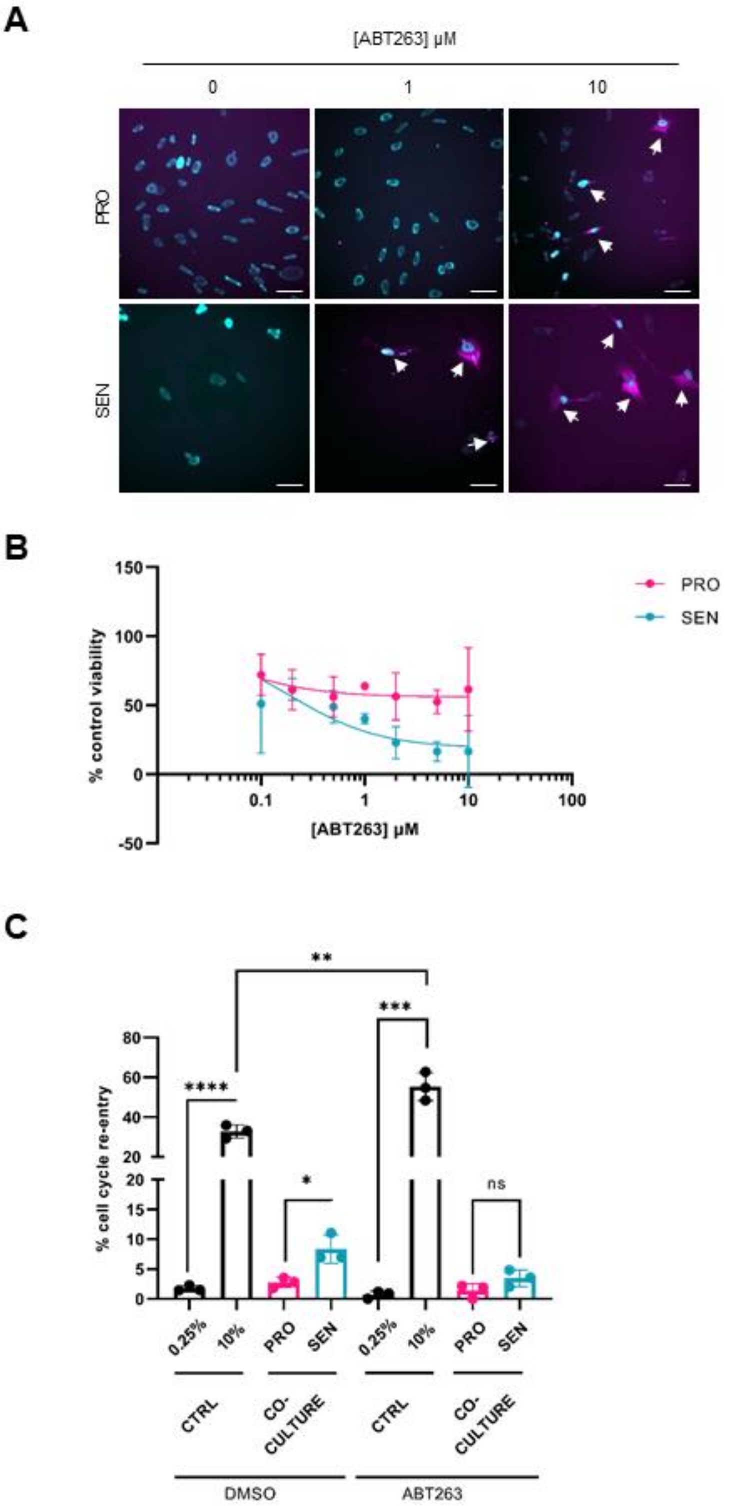
A
[ABT263] µM
0
1
10
PRO
SEN
B
% control viability
150
100
50
0
-50
0.1
1
10
100
[ABT263] µM
PRO
SEN
C
% cell cycle re-entry
80
60
40
20
20
15
10
5
0
****
**
***
*
ns
0.25%
10%
PRO
SEN
0.25%
10%
PRO
SEN
CTRL
CO-CULTURE
CTRL
CO-CULTURE
DMSO
ABT263

**Supplementary Figure 4. Pro-plasticity effects of senescent cells are abrogated by senolytic treatment**

**(A)** Representative images of cultured cells stained with NucBlue (DNA) and TO-PRO-3 iodide (apoptotic cells), 24h post-treatment with vehicle (DMSO) or 1 and 10 μM ABT-263. Arrows indicate inviable cells.

**(B)** Quantification of viable cells following fluorimetric assessment based on the cell metabolic indicator alamarBlue (n=3).

**(C)** Quantification of the proportion of myotube nuclei undergoing cell cycle re-entry, 72h post-treatment with the indicated conditions (DMSO vehicle control or ABT263 at 1 μM). Cultures were treated in 0.25% or 10% FCS control media, or in co-culture with proliferating or senescent cells in 0.25% FCS media. Cell cycle re-entry was quantified as the proportion of nuclei (stained using Hoechst) within myotubes (stained using α-MyHC) showing EdU incorporation. Two-tailed unpaired student's t-tests were used to compare data in **(C)** (*: $p<0.05$, **: $p<0.01$, ***:$p<0.001$, ****: $p<0.0001$, ns: not significant).

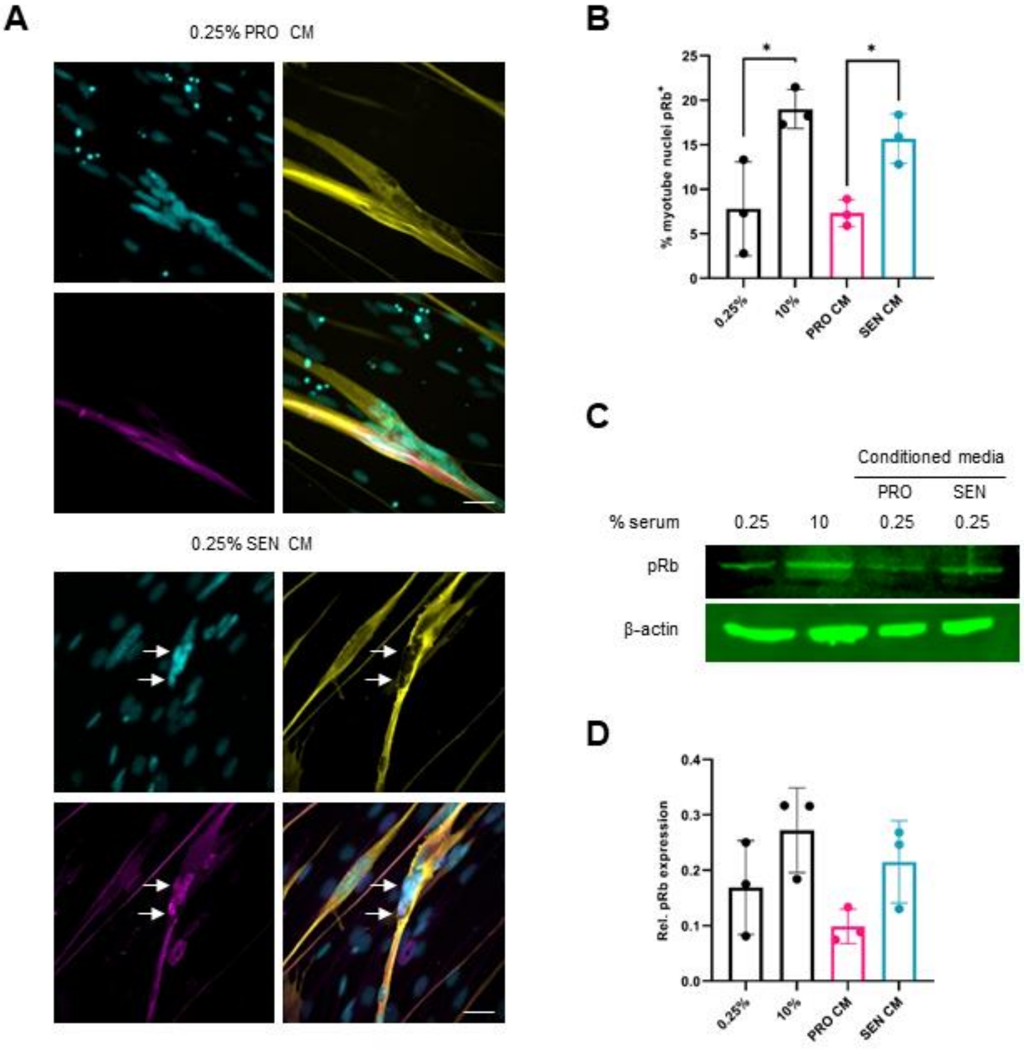
A
0.25% PRO CM
0.25% SEN CM
B
% myotube nuclei pRb+
0.25%
10%
PRO CM
SEN CM
C
Conditioned media
PRO
SEN
% serum
0.25
10
0.25
0.25
pRb
β-actin
D
Rel. pRb expression
0.25%
10%
PRO CM
SEN CM

**Supplementary Figure 5: Senescence-induced plasticity is accompanied by phosphorylation of Rb**

**(A)** Representative images of myotubes following immunostaining against MyHC (yellow), pRb (magenta) and Hoechst (cyan) labelling for the indicated conditions, 72hs post-treatment. White arrows indicate $EdU^+$ nuclei within myotubes.

**(B)**: Quantification of the proportion of myotube nuclei displaying pRb, as in **(B).** Statistical analysis performed using two-tailed unpaired student's t-test, *: $p<0.05$ (n=3).

**(C)**: Representative Western blot against pRb for the indicated conditions. β-actin was used as loading control.

**(D)**: Quantification of signal intensity. pRb band intensities were normalized against β-actin loading controls (n=3).

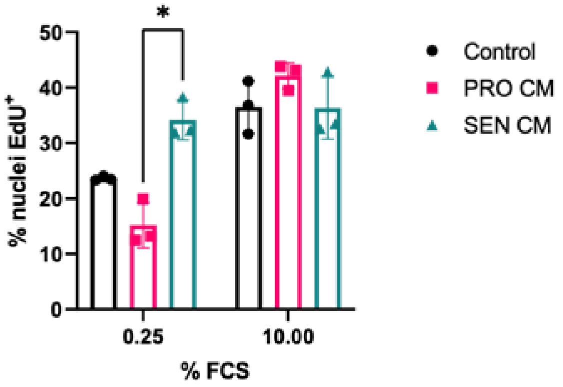

*
50
40
30
20
10
0
% nuclei EdU+
0.25
10.00
% FCS
Control
PRO CM
SEN CM

**Supplementary Figure 6: Senescence promotes cellular proliferation in a non-cell-autonomous manner**

A1 cells were exposed to fresh or 48 hour conditioned media containing 0.25% or 10% FCS for 72 hours before proliferation was assessed, as the % nuclei showing EdU incorporation (n=3, representative data from one of two independent experiments shown). ANOVA and post-hoc Tukey tests were used for statistical analysis (*: $p<0.05$, **: $p<0.01$, ns: not significant).

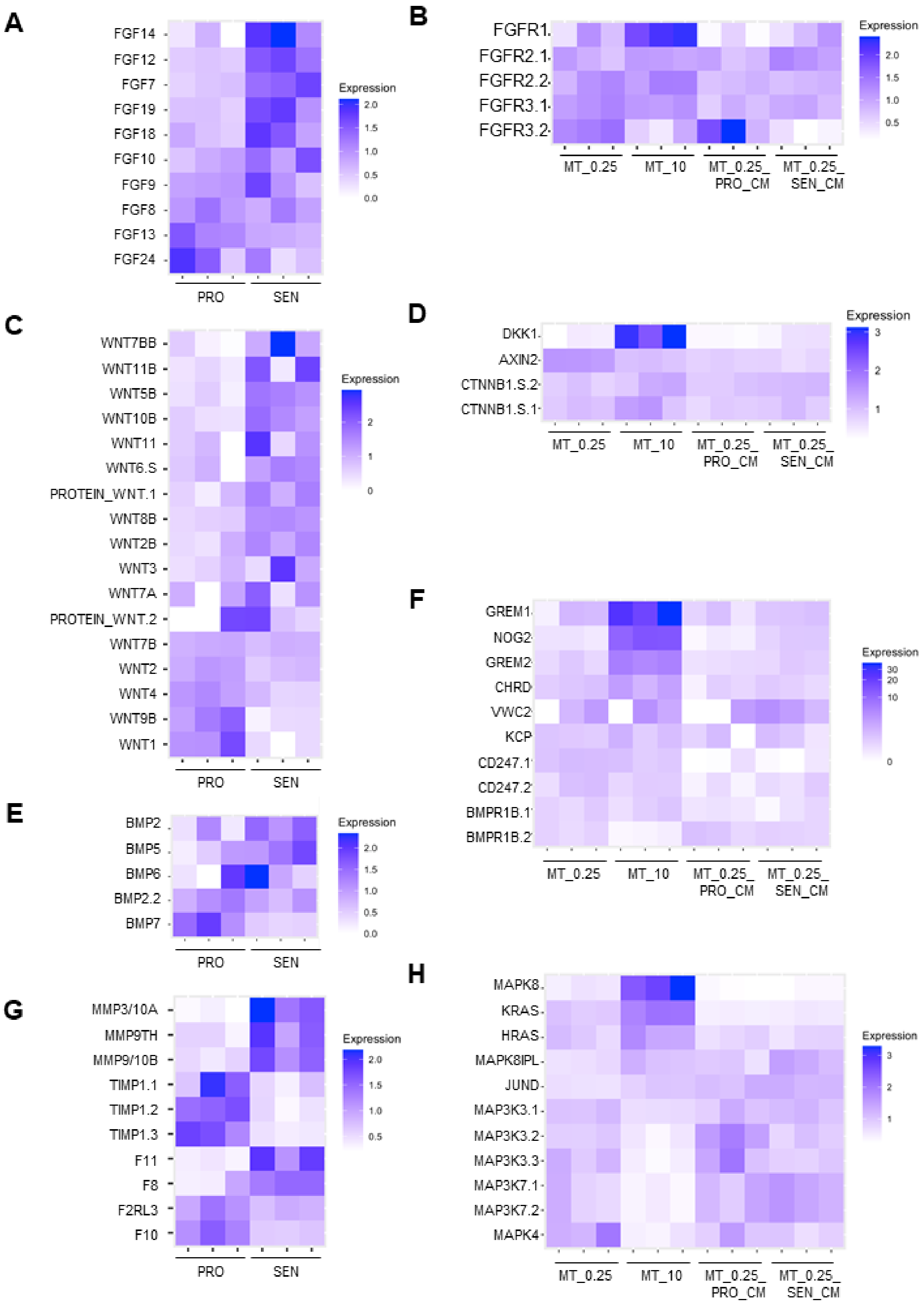
A
FGF14
FGF12
FGF7
FGF19
FGF18
FGF10
FGF9
FGF8
FGF13
FGF24
PRO
SEN
Expression
2.0
1.5
1.0
0.5
0.0
B
FGFR1
FGFR2.1
FGFR2.2
FGFR3.1
FGFR3.2
MT_0.25
MT_10
MT_0.25_ PRO_CM
MT_0.25_ SEN_CM
Expression
2.0
1.5
1.0
0.5
C
WNT7BB
WNT11B
WNT5B
WNT10B
WNT11
WNT6.S
PROTEIN_WNT.1
WNT8B
WNT2B
WNT3
WNT7A
PROTEIN_WNT.2
WNT7B
WNT2
WNT4
WNT9B
WNT1
PRO
SEN
Expression
2
1
0
D
DKK1
AXIN2
CTNNB1.S.2
CTNNB1.S.1
MT_0.25
MT_10
MT_0.25_ PRO_CM
MT_0.25_ SEN_CM
Expression
3
2
1
E
BMP2
BMP5
BMP6
BMP2.2
BMP7
PRO
SEN
Expression
2.0
1.5
1.0
0.5
0.0
F
GREM1
NOG2
GREM2
CHRD
VWC2
KCP
CD247.1
CD247.2
BMPR1B.1
BMPR1B.2
MT_0.25
MT_10
MT_0.25_ PRO_CM
MT_0.25_ SEN_CM
Expression
30
20
10
0
G
MMP3/10A
MMP9TH
MMP9/10B
TIMP1.1
TIMP1.2
TIMP1.3
F11
F8
F2RL3
F10
PRO
SEN
Expression
2.0
1.5
1.0
0.5
H
MAPK8
KRAS
HRAS
MAPK8IPL
JUND
MAP3K3.1
MAP3K3.2
MAP3K3.3
MAP3K7.1
MAP3K7.2
MAPK4
MT_0.25
MT_10
MT_0.25_ PRO_CM
MT_0.25_ SEN_CM
Expression
3
2
1

**Supplementary Figure 7. Candidate pathways involved in mediating senescence-induced plasticity*.***

**(A-H)** heatmaps depicting transcriptional changes between proliferating vs. senescent cells **(A, C, E, G)** or between differentiated and dedifferentiated myotubes upon serum or senescent CM exposure **(B, D, F, H)**. Ligands and downstream signalling transcripts are depicted for the FGF pathway **(A, B)**, for Wnt signalling **(C, D)** for BMP signalling **(E, F)** and for protease expression **(G)** and ERK signalling **(H)**. Transcript expression for each replicate is normalized relative to the mean reads per million transcripts across all samples groups in each heatmap.

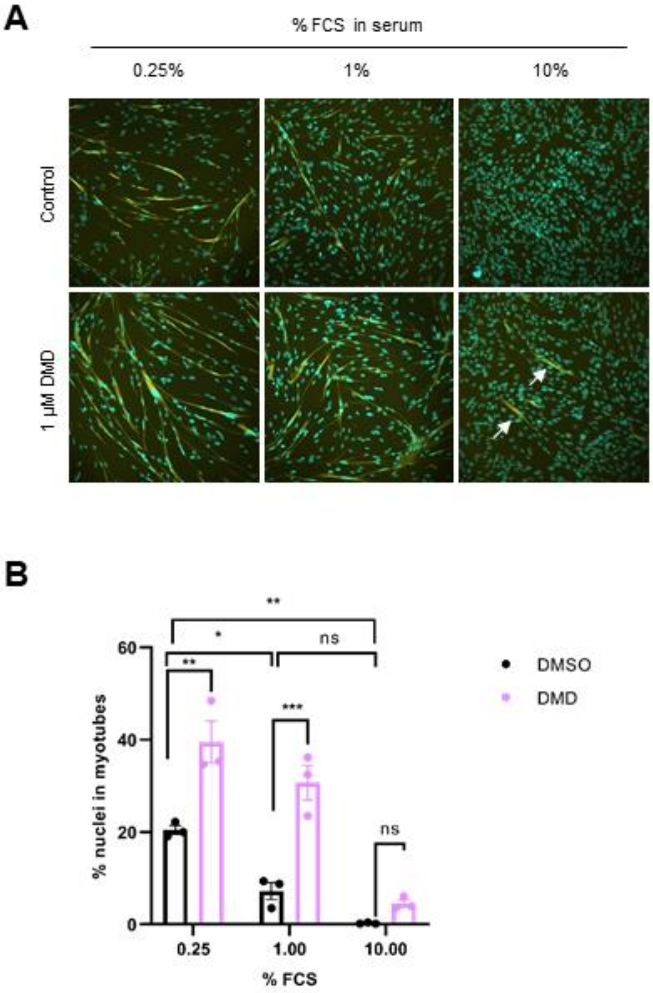
A
% FCS in serum
0.25%
1%
10%
Control
1 µM DMD
B
% nuclei in myotubes
60
40
20
0
0.25
1.00
10.00
% FCS
**
*
ns
***
DMSO
DMD

**Supplementary Figure 8. BMP inhibition promotes myogenesis.**

**(A, B)** A1 cells were seeded at high confluence and left to bed down before exposure to 1 μM dorsomorphin (DMD) or vehicle only (DMSO) in 0.25%, 1% or 10% FCS-supplemented media for 5 days. Cells were then fixed and immunostained with α-MyHC and counter-stained with Hoechst. Representative images shown in **(A)**, quantification of % nuclei in myotubes in **(B)**. White arrows indicate myotube formation in 10% serum upon DMD treatment. Two-way ANOVA and post-hoc Tukey tests were used for statistical analysis (*: $p<0.05$, **: $p<0.01$, ***: $p<0.001$, ns: not significant). Representative data shown from 2 independent experiments (n=3 technical replicates).

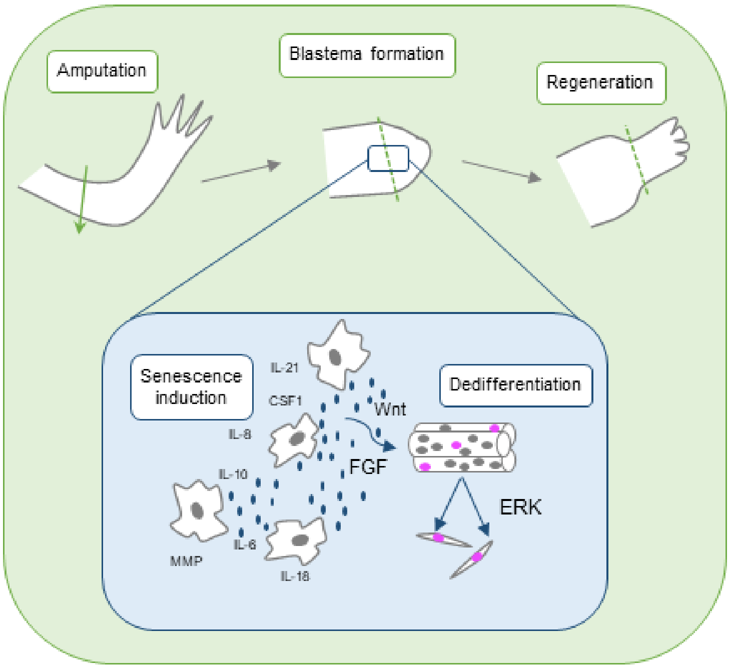
Amputation
Blastema formation
Regeneration
Senescence induction
Dedifferentiation
IL-21
CSF1
IL-8
IL-10
MMP
IL-6
IL-18
Wnt
FGF
ERK

**Supplementary Figure 9. Mechanism of promotion of dedifferentiation by senescent cells during newt limb regeneration.**

Schematic depiction of the proposed mechanism for how senescence induction enhances limb regeneration. Senescent cells, present over a short time-window during early regeneration stages, promote muscle dedifferentiation in a non-cell autonomous manner. This results in activation of the FGF-ERK signalling axis in differentiated muscle to form myogenic progenitor cells. Following many rounds of proliferation within the blastema, these progenitors redifferentiate, in a process possibly involving BMP inhibition, to accomplish muscle regeneration.

**Table S1. Inhibitors used in this study**

| Inhibitor | Target | Dose used (μM) |
|---|---|---|
| ABT263 | BCL2 (senolytic) | 1 |
| Dorsomorphin | BMP antagonist | 1 |
| U0126 | MEK1/2 (ERK) | 10 |
| PD173074 | FGFR1 | 1 |
| AZD4547 | FGFR1,2,3 | 5 |
| C59 | Wnt | 1 |
| AEBSF | Serine protease | 20 |
| GM6001 | Broad spectrum MMP | 2 |
| Etoposide | Topoisomerase II | 20 |
| Nutlin-3a | p53/MDM2 | 1 |

**Table S2. Antibodies used in this study.**

| Target | Supplier and code | Application | Dilution |
|---|---|---|---|
| **Primary** | | | |
| α -myosin heavy chain | Custom | Immunofluorescence | 1:1000 |
| | | Western blotting | 1:2000 |
| α-phospho-Rb | Cell signalling (9308) | Immunofluorescence | 1:200 |
| | | Western blotting | 1:500 |
| α-GFP | Abcam (ab6673) | Immunofluorescence | 1:1000 |
| **Secondary** | | | |
| α-mouse 680/800<br>α -rabbit 680/800 | Licor | Western blotting | 1:2000 |
| Alexafluor α-mouse/rabbit 488/594/647 | Invitrogen | Immunofluorescence | 1:1000 |

### Other Supplementary Materials for this manuscript include the following:

**Data S1. DGE analysis for all conditions described in Figure 3**

**Data S2. List of genes corresponding to the GO pathway enrichment analysis corresponding to Figure 3 and adjusted p-values.**
